# Decoding the relative contributions of extrinsic and intrinsic mechanisms in mediating heterogeneous spiking activities of sensory neurons in vivo using computational modeling

**DOI:** 10.1101/2023.01.03.521866

**Authors:** Amin Akhshi, Myriah Haggard, Mariana M. Marquez, Saeed Farjami, Maurice J. Chacron, Anmar Khadra

**Author notes:** Authors contributed equally (Maurice J. Chacron, Anmar Khadra).

## Abstract

Neurons ubiquitously display heterogeneities in spiking activity even within a given cell type. To date, the relative contributions of extrinsic mechanisms (e.g., synaptic bombardment) and intrinsic mechanisms (e.g., conductances, cell morphology) towards determining spiking activity remain poorly understood. Here we address this important question using a novel approach that combines biophysical techniques, in which extracellular in vivo recordings of electrosensory pyramidal cells within weakly electric fish, are combined with computational modeling. Specifically, by varying parameters, a conductance-based computational model successfully reproduced the highly heterogeneous spiking activities seen experimentally. Model parameters that varied the most were then used to gauge the relative contributions of extrinsic vs. intrinsic mechanisms. Overall, extrinsic synaptic input was predicted to be the main factor accounting for spiking heterogeneities. We tested this prediction experimentally by performing two different manipulations: i) pharmacologically inactivating feedback; ii) applying the neuromodulator serotonin. Our model predicted that feedback inactivation should reduce while serotonin application should increase spiking heterogeneities. Experiments corroborated these predictions. Importantly, for serotonin application, increased heterogeneity occurred despite a strong reduction in intrinsic membrane conductance, further demonstrating that extrinsic synaptic input is the primary determinant of spiking heterogeneities in vivo. Taken together, our results demonstrate that devising a computational model to capture spiking heterogeneities in vivo and assessing which parameters are responsible can successfully determine the relative contributions of extrinsic vs. intrinsic inputs. We expect this approach to be generalizable to other systems and species.

## Introduction

It has been observed ubiquitously across brain areas that sensory neurons, even if they encode the same stimulus property, display large heterogeneities in their spiking activities (1–5). While growing evidence suggests that heterogeneous firing patterns can be beneficial, by effectively increasing their coding range compared to homogeneous neural activities as well as promoting more robust learning in neural circuits (6–13), how different mechanisms contribute to generating them is not yet completely understood.

While spiking heterogeneities can be the result of multiple factors, including morphological (14, 15), molecular (16–18), and electrophysiological (19) differences, their relative contributions remain obscure. This is because not all of these necessarily give rise to spiking heterogeneity. Indeed, previous studies have demonstrated that neural networks can have very similar spiking activities despite large differences in their synaptic connection strength and/or the presence and expression of different membrane conductances (20–22) (see (23) for review). One potent approach to understanding how different factors contribute to spiking heterogeneities is computational modeling. Specifically, recent approaches have shown that computational models can be successfully used to link electrophysiological properties with different factors, including gene expression, to correctly reproduce spiking heterogeneities seen in vitro (24, 25). However, whether such an approach can successfully explain such relationships in vivo, where neurons are in a high-conductance state due to massive synaptic bombardment (26) remains unknown to date.

The firing patterns exhibited by sensory pyramidal cells within the electrosensory lateral line lobe (ELL) of gymnotiform wave-type weakly electric fish in vivo constitute prototypical examples of neural activities displaying high level of spiking heterogeneity (27–29) (see (30–34) for review). ELL pyramidal cells receive feedforward inputs from electroreceptor afferents located on the skin surface and, as the sole output neurons of the ELL, project to higher brain regions that mediate perception and behavior (35, 36). Previous studies have shown that there are large levels of heterogeneity within the ELL pyramidal cell population in terms of cellular morphology, expression of intrinsic receptors, as well as the amount of extrinsic synaptic input (see (30) for review). Interestingly, these ELL pyramidal cells are organized into columns consisting of three ON- and three OFF-type cells, labeled “superficial”, “intermediate”, and “deep” based on the location of their somata within the pyramidal cell layer (30). Superficial, intermediate, and deep pyramidal cells differ in their morphologies, with superficial cells displaying the largest apical dendrites and deep cells displaying the smallest (27, 37, 38). Previous studies have also shown differential expression of various intrinsic receptors, with deep pyramidal cells showing the lowest and superficial pyramidal cells the highest expression levels of membrane conductances, such as small-conductance calcium activated potassium (SK) channels (39), NMDA receptors (40), as well as IP3 receptors (IP3Rs) (see (30) for review).

Finally, ELL pyramidal cells also receive differential amounts of descending synaptic input (i.e., feedback) from higher brain regions (41), with superficial cells receiving the largest amount and deep cells the smallest (38). Such feedback can strongly alter neural responses to feedforward input through gain control and cancellation of redundant stimuli (29, 42–44), as well as generate responses to sensory input arising during specific behavioral contexts (45–48). ELL pyramidal cells also receive differential amounts of neuromodulatory input including serotonergic fibers from the raphe nuclei (49, 50). Serotonin application downregulates SK channels, thereby increasing neural excitability (50), as well as responses to sensory input (51–53) (see (33, 54, 55) for review). How these different factors contribute to mediating the observed large heterogeneities in spiking activities observed in vivo remains unclear. Here we used a combination of computational modeling and in vivo electrophysiological recordings from ELL pyramidal cells to investigate the relative contributions of feedback synaptic input, intrinsic membrane conductances, and cellular morphology towards determining differences in spiking activity.

## Results

### ELL pyramidal cell spiking activities are highly heterogeneous in vivo

The goal of the current study was to gain understanding as to the relative contributions of multiple mechanisms such as synaptic input, intrinsic membrane conductances, and cellular morphology towards generating heterogeneities in spiking activities (Fig. 1A). To do so, we recorded the spiking activities of n=243 ELL pyramidal cells across N=30 fish in vivo. In most cases, recordings were achieved using high-density arrays (i.e., Neuropixels probes) into the brain (Fig. 1B) (see Methods). ELL pyramidal cells receive feedforward input from peripheral electroreceptor afferents and project indirectly to higher brain areas mediating perception and behavior via the torus semicircularis (TS) (Fig. 1C). ELL pyramidal cells also receive a large amount of feedback input from the nucleus praeminentialis (nP) (Fig. 1C). Overall, examination of spiking activities of these cells revealed that they were highly heterogeneous (Fig. 1D). Indeed, large differences were seen in their mean firing rate which ranged between 3 Hz and 63 Hz, consistent with previous studies (27, 28, 56). Neural activities also strongly differed in terms of the tendency to fire packets of action potentials followed by quiescence (i.e., bursts) (e.g., compare cells i and iv in Fig. 1D). We used the interspike interval (ISI) distributions, that quantify the time between consecutive action potentials (APs) within each recording, to characterize spiking heterogeneities across the ELL pyramidal cell population. This is because previous studies have shown that such distributions can account for differences in mean firing rate as well as burst firing between cells (27, 57). As expected, ISI distributions were highly heterogeneous (Fig. 1E).

**Fig. 1.**
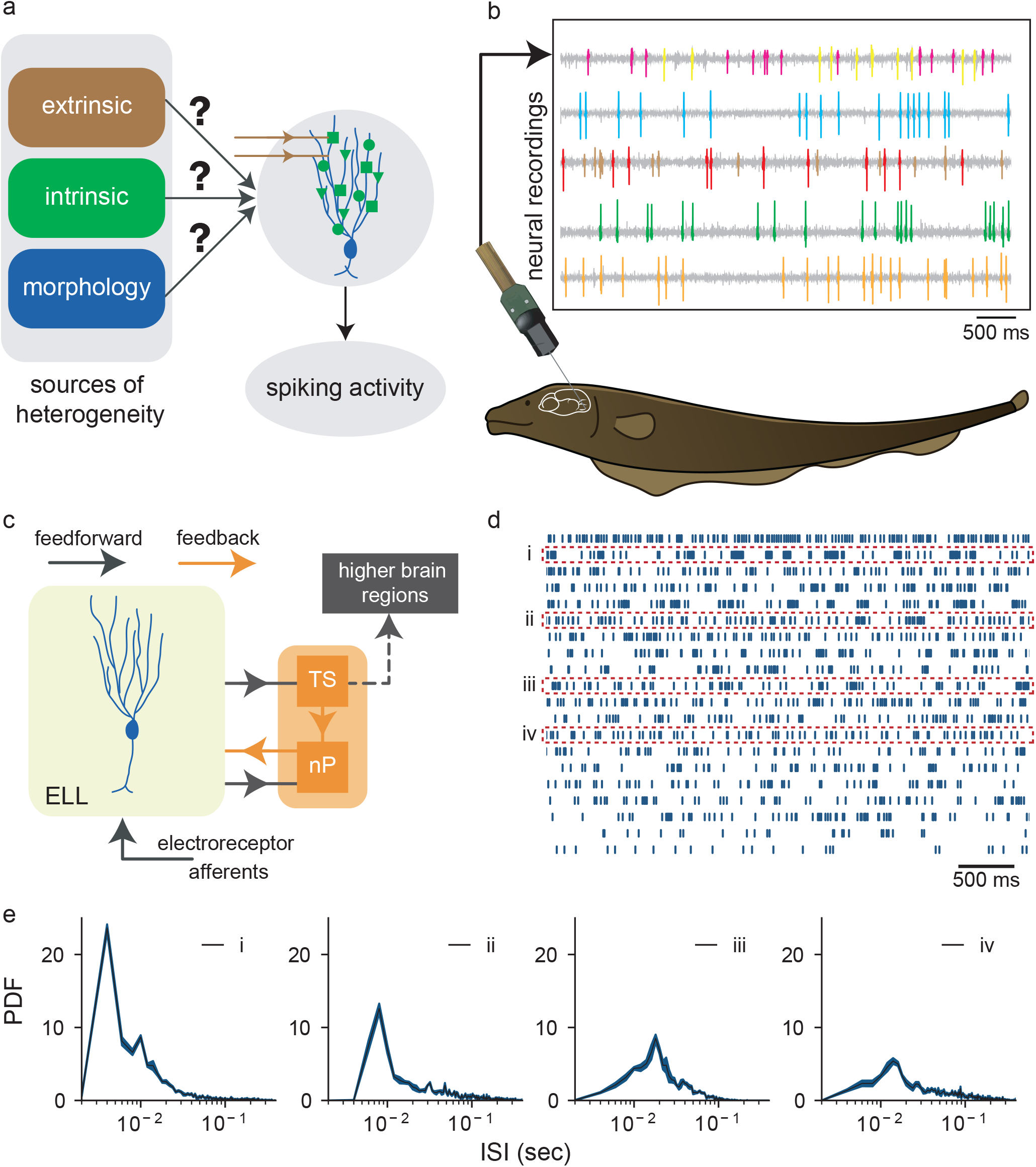
ELL pyramidal cells display heterogeneous spiking activities in vivo. **A)** Schematic showing how different factors can contribute to heterogeneous spiking activities. **B)** An outline of the recording setup. Extracellular recordings from ELL pyramidal cells were made using high-density arrays and the spikes were sorted (top left). **C)** Simplified diagram of feedforward (black) and feedback (orange) electrosensory pathways. Electroreceptor afferents project to pyramidal cells within the electrosensory lateral line lobe (ELL), which in turn project directly onto the midbrain torus semicircularis (TS) and indirectly to higher brain areas. ELL pyramidal cells also receive feedback input from the nucleus praeminentialis (nP). **D)** Raster plot showing the spiking activities of 20 representative ELL pyramidal cells sorted from highest to lowest firing rate from top to bottom. These activities were not recorded simultaneously. **E)** Interspike interval (ISI) probability densities from four example ELL pyramidal cells whose spiking activities are indicted in panel D.

### Varying parameter values in a conductance-based mathematical model gives rise to spiking activities that closely mimic those seen experimentally

In order to understand the relative contributions of extrinsic synaptic input, intrinsic membrane conductances, and cellular morphology towards determining spiking activity in vivo, we built a new computational model of ELL pyramidal cell activity that was in part inspired by previous studies (58, 59), but also included calcium mobilization as well as synaptic bombardment (Fig. 2A, see Methods). The model is comprised of two compartments (soma and dendrite) that are coupled through a resistance, with each compartment containing different conductances (Fig. 2A, left). Calcium dynamics were implemented in the dendritic compartment by including calcium fluxes across both the cell and ER membranes (Fig. 2A, right). When simulating the model, we obtained time courses for the somatic and dendritic membrane potentials as well as physiologically realistic cytosolic and ER calcium transients in the dendrite (Supplementary Fig. 1).

**Fig. 2.**
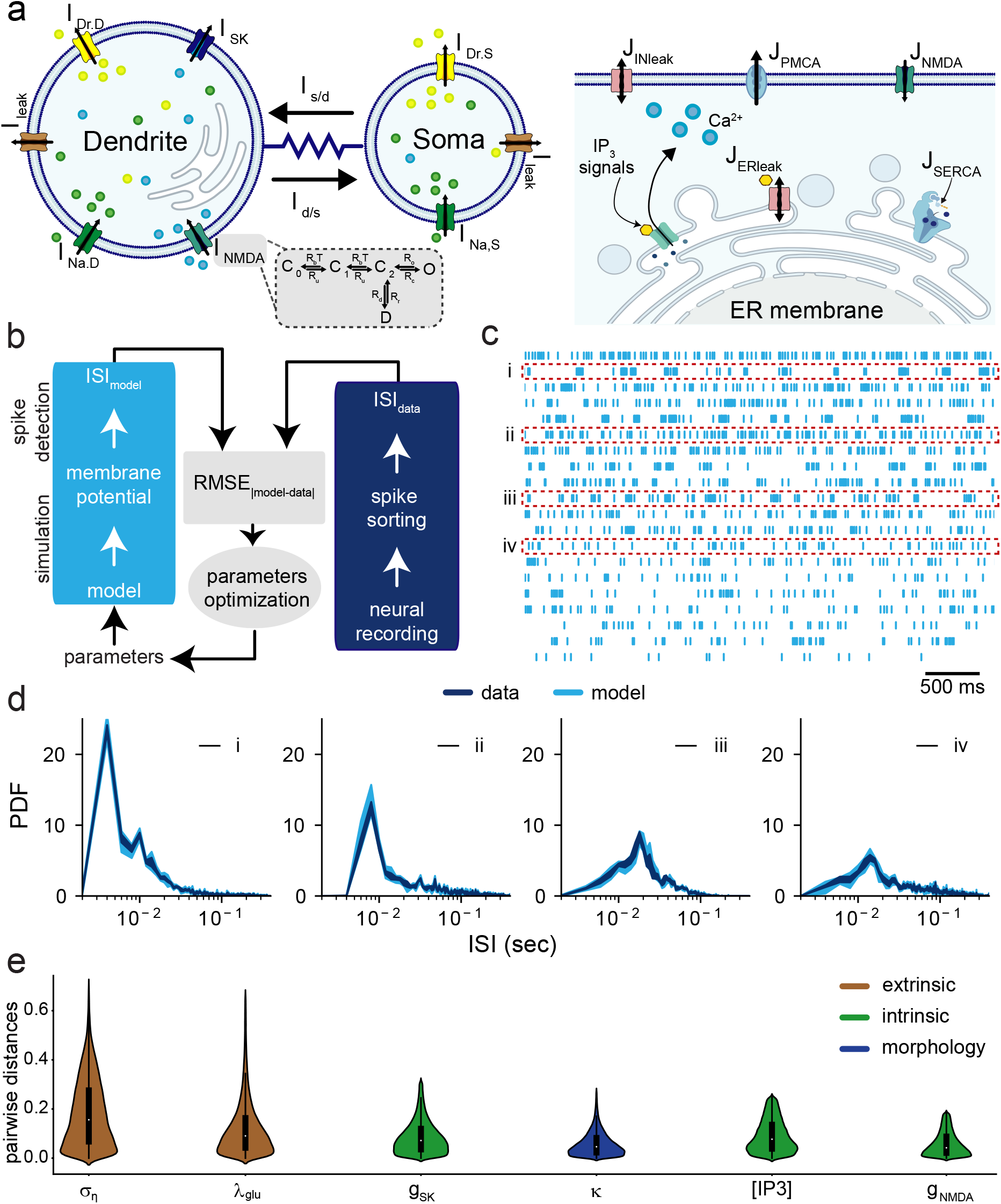
A conductance-based computational model correctly reproduces spiking heterogeneities seen in vivo across ELL pyramidal cells and predicts that they are primarily due to differences in extrinsic synaptic input. **A)** Schematic of the model highlighting the two-compartments of the cell: soma and dendritic tree with the various ionic currents expressed in each compartment, including fast sodium (I_Na,i_), delayed rectifier potassium (I_K,i_), small-conductance potassium (I_SK_), NMDA (I_NMDA_) and leak (I_leak_) currents in the soma (i = S) and dendrite (i = D) (top left). The kinetics of NMDA receptors are governed by a Markov model (gray box) consisting of three closed states (C_0_,C_1_,C_2_), one conducting open state (O) and one desensitized state (D). Within the dendritic compartment, calcium dynamics are implemented using the flux-balance formalism. Shown are all fluxes involved in regulating calcium mobilization across the cell and ER membranes in the dendrite. These fluxes include calcium entry through NMDA receptors (J_NMDA_) and IP3Rs (J_IP3R_), calcium efflux through SERCA (J_SERCA_) and PMCA (J_PMCA_) pumps, and leak across both membranes (top right). **B)** Schematic showing the algorithm used to fit the model to each ELL pyramidal cell that was recorded from. A maximum likelihood estimation was used to find model parameters for which the root mean square error (RMSE_|model-data|_) between the ISI distributions obtained from the model (ISI_model_) and experimental data (ISI_data_) was minimized. **C)** Spiking activities obtained from the model for each ELL pyramidal cell whose activity was shown in Fig. 1D. **D)** ISI distributions obtained from the model (light blue) and experimentally (dark blue) for the same four example ELL pyramidal cells shown in Fig. 1E. **E)** Violin plots showing the distributions of pairwise distances for the six model parameters that needed to be varied the most in order to correctly reproduce spiking heterogeneities seen experimentally sorted in descending order. Overall, model paramate *σ_η_* (the standard deviation of extrinsic synaptic input) and λ_glu_ (the mean extrinsic synaptic input rate) needed to be varied the most, followed by g_SK_ (the SK channel maximum conductance), *κ* (the somatic-to-dendritic area ratio), [IP3] (the concentration of IP3) and g_NMDA_(the NMDA receptor maximum conductance).

We next tested whether it was possible to find parameter values for our model that would reproduce the heterogeneous firing activities of ELL pyramidal cells seen in vivo. Specifically, we asked whether the ISI distribution obtained from the model can be matched to those seen experimentally for each single neuron that was recorded from. To do so, we used a maximum likelihood algorithm to minimize the error between the ISI distributions obtained from the model and experimentally (Fig. 2B). The algorithm was initiated from a set of parameter values obtained from previous studies (58, 59) and the error between the model and data, quantified using root mean squared error (RMSE), was minimized with maximum likelihood (see Methods). Our results showed that the model, under different parameter regimes, could give rise to spiking activities that closely resembled those seen experimentally (compare Fig. 2C with Fig. 1D). Indeed, ISI distributions obtained from the model closely matched those seen experimentally (Fig. 2D) both in terms of mean firing rate but also in the tendency to fire bursts of action potentials. Taken together, these results demonstrate that our model successfully reproduced the heterogeneous firing patterns observed experimentally in vivo across the ELL pyramidal cell populations.

In order to gain further insight as to the relative contributions of extrinsic synaptic input, intrinsic properties (such as membrane conductances and calcium mobilization) and cellular morphology, we investigated which model parameters needed to be varied the most in order to account for spiking heterogeneities. Specifically, we looked at the distributions of normalized pairwise distances between parameter values for which the model fitted experimental data from a given cell (see Methods). Figure 2E shows the distributions of pairwise distances for the six parameters with the greatest ranges sorted in descending order. Overall, the two model parameters that needed to be varied the most, *σ_η_* and λ_glu_, both directly influenced extrinsic synaptic input in terms of intensity and mean firing rate level, respectively. Of the other four parameters, three were associated with intrinsic mechanisms, including SK channel maximum conductance (g_SK_), the concentration of IP3 ([IP3]) and the NMDA receptor maximum conductance (g_NMDA_), while the other was associated with cellular morphology (i.e., the somatic-to-dendritic area ratio *κ*). Thus, our model makes the important prediction that spiking heterogeneities seen in vivo across the ELL pyramidal cell population are primarily due to differences in extrinsic synaptic input and, to a lesser extent, differences in intrinsic properties and morphology.

### Pharmacological inactivation of feedback pathways reduces spiking heterogeneities across the ELL pyramidal cell population

So far, our results show that our computational model is able to not only reproduce the heterogeneous spiking activities displayed by ELL pyramidal cells in vivo, but also to make predictions that heterogeneities are due to distinct ELL pyramidal cells receiving different levels of extrinsic synaptic input. In order to test this prediction, we pharmacologically inactivated feedback input onto ELL pyramidal cells by injecting the sodium channel antagonist lidocaine within nP (Fig. 3A). Such a manipulation is expected to significantly reduce the amount of synaptic input received by ELL pyramidal cells, which according to our modeling predictions should render their spiking activities less heterogeneous. Figure 3B shows the spiking activities of the same four ELL pyramidal cells before (i.e., under control conditions; dark blue) and after (orange) pharmacological inactivation. As predicted, heterogeneities between cells are reduced, with the four cells exhibiting spiking activities that closely resemble one another after inactivation (Fig. 3B, compare dark blue and orange); this is in part due to the fact that tendency to fire bursts of action potentials was reduced overall (Fig. 3B) as reflected by changes in the ISI distribution (sup Fig. 2A). In order to visualize the effects of pharmacological inactivation, we used principal component analysis to reduce the dimensionality of our data (see Methods); with this framework, each neuron was represented by one single datapoint in a space spanned by the first three principal components, which together accounted for 82% of the variance. Figure 3C shows plots of our data before (i.e., “control”; left panel) and after (i.e., “block”; right panel) pharmacological inactivation. It can be seen that the datapoints occupy a smaller volume in space after inactivation as compared to control conditions (Fig. 3C, compare orange and dark blue). To quantify this change, we computed both the pairwise distance as well as the volume occupied by the datapoints (see Methods). In agreement with our latter observation, the ratios of values obtained for both pairwise distance and volume after inactivation to those obtained before inactivation were significantly less than unity (Fig. 3D; volume: **p = 0.00002, two-sample t-test; pairwise distance: *p = 0.0001, two-sample t-test). These results thus highlight that pharmacological inactivation of feedback pathways leads to reduced heterogeneity in the spiking activities of ELL pyramidal cells in vivo, consistent with our modeling predictions.

**Figure 3:**
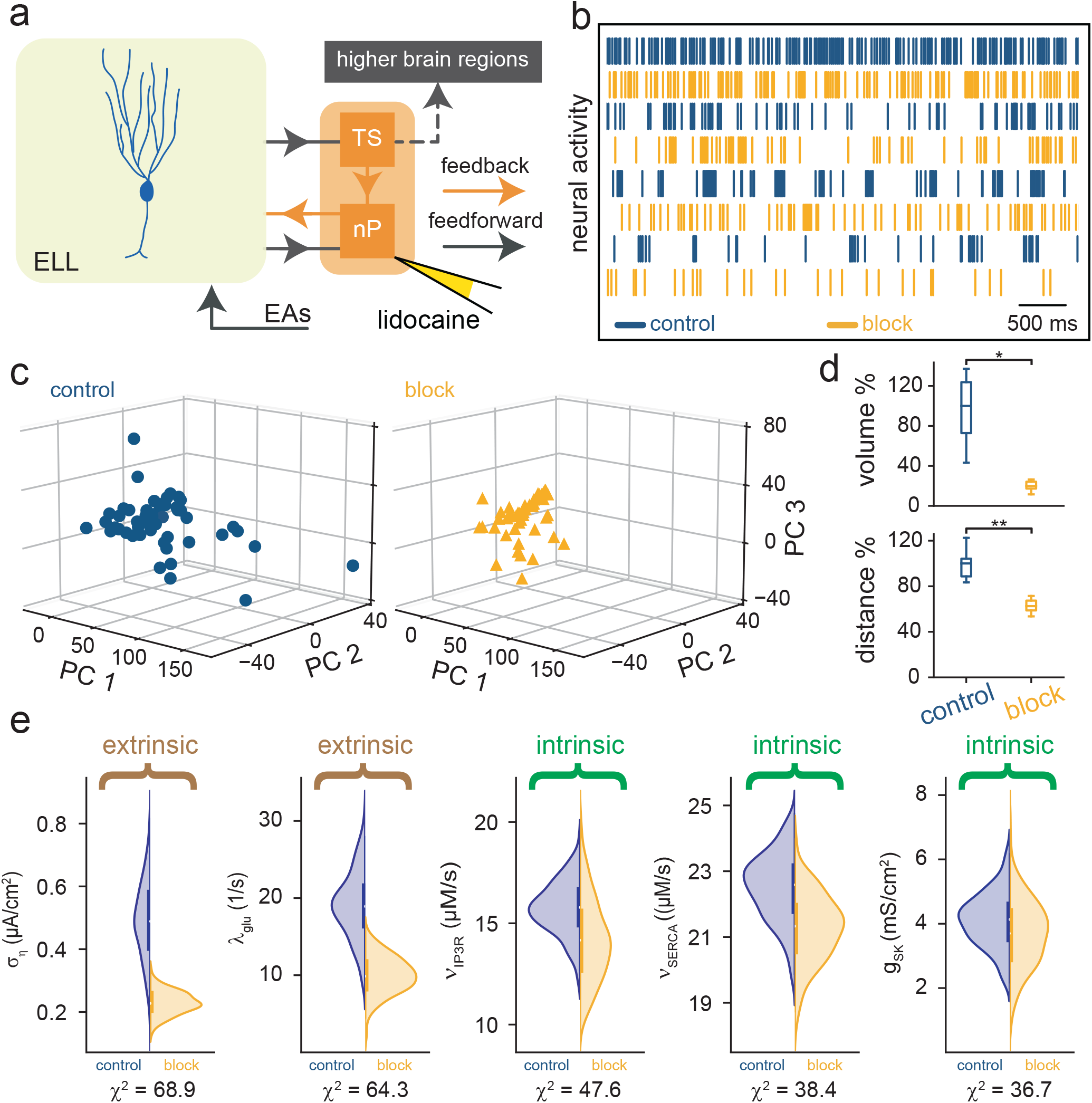
Pharmacological inactivation of feedback pathways reduces spiking heterogeneities within the ELL pyramidal cell population. **A)** Simplified diagram of feedforward (black) and feedback (orange) electrosensory pathways. Pharmacological inactivation of feedback pathways was achieved by injecting the sodium channel antagonist lidocaine within nP bilaterally (see Methods). **B)** Spiking activities of four example ELL pyramidal cells before (dark blue) and after (orange) inactivation of feedback pathways. **C)** Plots of our dataset using the first three principal components (i.e., PC 1, PC 2, and PC 3) that together account for 82%of the variance before (left) and after (right) pharmacological inactivation, where each datapoint represents a neuron. Notice how the datapoints occupy a significantly smaller volume after inactivation, thereby implying lower pairwise distances on average. **D)** Box plots showing that the block-to-control volume (top) and pairwise distance (bottom) ratios were significantly decreased (volume: *p = 0.00002, two-sample t-test; pairwise distance: **p = 0.0002, two-sample t-test). **E)** Violin plots showing distributions of pairwise distances before (blue, left) and after (orange, right) pharmacological inactivation for the top six model parameters that needed to be varied significantly in order to account for changes in spiking activity seen experimentally. Overall, the values of the extrinsic parameters *σ_η_* (the standard deviation of extrinsic synaptic input) and λ_glu_ (the mean extrinsic synaptic input rate) needed to be reduced the most, while the values of other intrinsic parameters such as *ν*_IP3R_ (the IP3R maximum flux rate), *ν*_SERCA_ (SERCA maximum flux rate), and g_SK_ (the SK channel maximum conductance) needed to be reduced to a lesser extent.

To further confirm these results, we fitted our model to the experimental data obtained from ELL pyramidal cells after pharmacological inactivation using the same methodology described above (Fig. 2B). As before, ISI distributions obtained from the model closely matched those seen experimentally (sup Fig. 2B). We then determined which model parameters varied the most in order to account for experimentally observed changes in spiking activity. Since experimental data were obtained from the same neurons before and after pharmacological inactivation, we compared the distributions of parameter values in our model when fitted to experimental data under both conditions (see Methods). Figure 3E shows distributions of values obtained when fitting experimental data before (blue) and after (orange) pharmacological inactivation for the six parameters for which the distributions significantly differed from one another, with significance quantified by the *χ*^2^ value (note that higher values of *χ*^2^ imply a greater degree of significance). Overall, our results showed that, in order to correctly account for changes in spiking activity induced by pharmacological inactivation, the values of the two parameters *σ_η_* and λ_glu_, which directly influence extrinsic synaptic input, needed to be strongly reduced. This is to be expected since our previous findings indicated that pharmacological inactivation should strongly reduce the overall level of synaptic input received by ELL pyramidal cells. Our results also showed that other parameters associated with intrinsic properties, including *ν*_IP3R_ (i.e., the IP3R maximum flux rate), *ν*_SERCA_ (i.e., the maximum flux rate of sarco/endoplasmic reticulum calcium ATPases (SERCA)) and g_SK_ (i.e., the SK maximum channel conductance), needed to be reduced but to a lesser extent. Interestingly, such changes are also expected given that blocking synaptic input alters calcium influx and mobilization in the cell, which will directly affect SK channels that are sensitive to the internal calcium concentration.

### Exogenous application of serotonin increases spiking heterogeneities across the ELL pyramidal cell population

As a further test of our modeling prediction that spiking heterogeneities across the ELL pyramidal cell population are primarily due to differences in extrinsic synaptic input, we applied the neuromodulator serotonin (Fig. 4A; see Methods). Previous studies have shown that serotonin application strongly alters the spiking activities of ELL pyramidal cells through increases in the mean firing rate and the tendency to fire bursts of action potentials, which occurs in part because of inhibition of SK channels (50–53, 60). Assuming that serotonin application does not alter any other mechanism, one would thus predict (based on our previous results in Fig. 2E) that, because the SK channel conductance should be reduced after serotonin application, spiking heterogeneities should be reduced. Figure 4B shows the spiking activities of the same four ELL pyramidal cells before (dark blue) and after (red) serotonin application. Overall, consistent with previous results, there was an overall increase in the mean firing rate and burst firing upon serotonin application (Fig. 4B, compare dark blue and red; sup Fig. 3A). Interestingly however, spiking activities were more heterogeneous from one another after serotonin application (Fig. 4B, compare red and dark blue). Plotting the data in the space spanned by the first three principal components revealed that datapoints did occupy a larger volume (Fig. 4C, compare red and dark blue). Indeed, for both pairwise distance and volume, the ratios of values obtained after inactivation to those obtained before inactivation were both significantly greater than unity (Fig. 4D; volume: *p = 0.00002, two-sample t-test; pairwise distance: **p = 0.0002, two-sample t-test). These experimental data thus shows that serotonin application actually increases spiking heterogeneities across the ELL pyramidal cell population. In order to gain better understanding as to why serotonin application actually increases spiking heterogeneities rather than decreasing it (in view of the fact that serotonin reduces SK channel maximum conductance), we fitted our model to experimental data obtained after serotonin application. Overall, our results showed a close match between ISI distributions obtained from our model and those obtained experimentally (sup Fig. 3B). We then compared the distributions of parameter values in our model when fitted to experimental data under both conditions in a manner similar to what was done before for pharmacological inactivation (see Methods). Figure 4E shows distributions of values obtained when fitting experimental data before (blue) and after (red) serotonin for the four parameters for which the distributions significantly differed from one another, with significance again quantified using the *χ*^2^ value. Overall, the SK channel maximum conductance needed to be reduced in order for our model to correctly reproduce experimental data obtained after serotonin application (Fig. 4E, left), in agreement with experimental data showing that SK currents are inhibited by serotonin (50). However, our results also showed that the intensity of extrinsic synaptic input, as parametrized by *σ_η_*, also needed to be increased (Fig. 4E, center left). The values of two other parameters, the NMDA receptor maximum conductance g_NMDA_ and the extracellular magnesium concentration [Mg^2+^]_o_ also needed to be reduced but to a lesser extent (Fig. 4E, center right and right). Thus, our model provides an explanation as to why serotonin application increases spiking heterogeneities. Specifically, the increase in heterogeneity from increased intensity of extrinsic synaptic input trumps the decrease in heterogeneity from decreased SK channel conductance, thereby leading to an overall increase in heterogeneity.

**Fig. 4.**
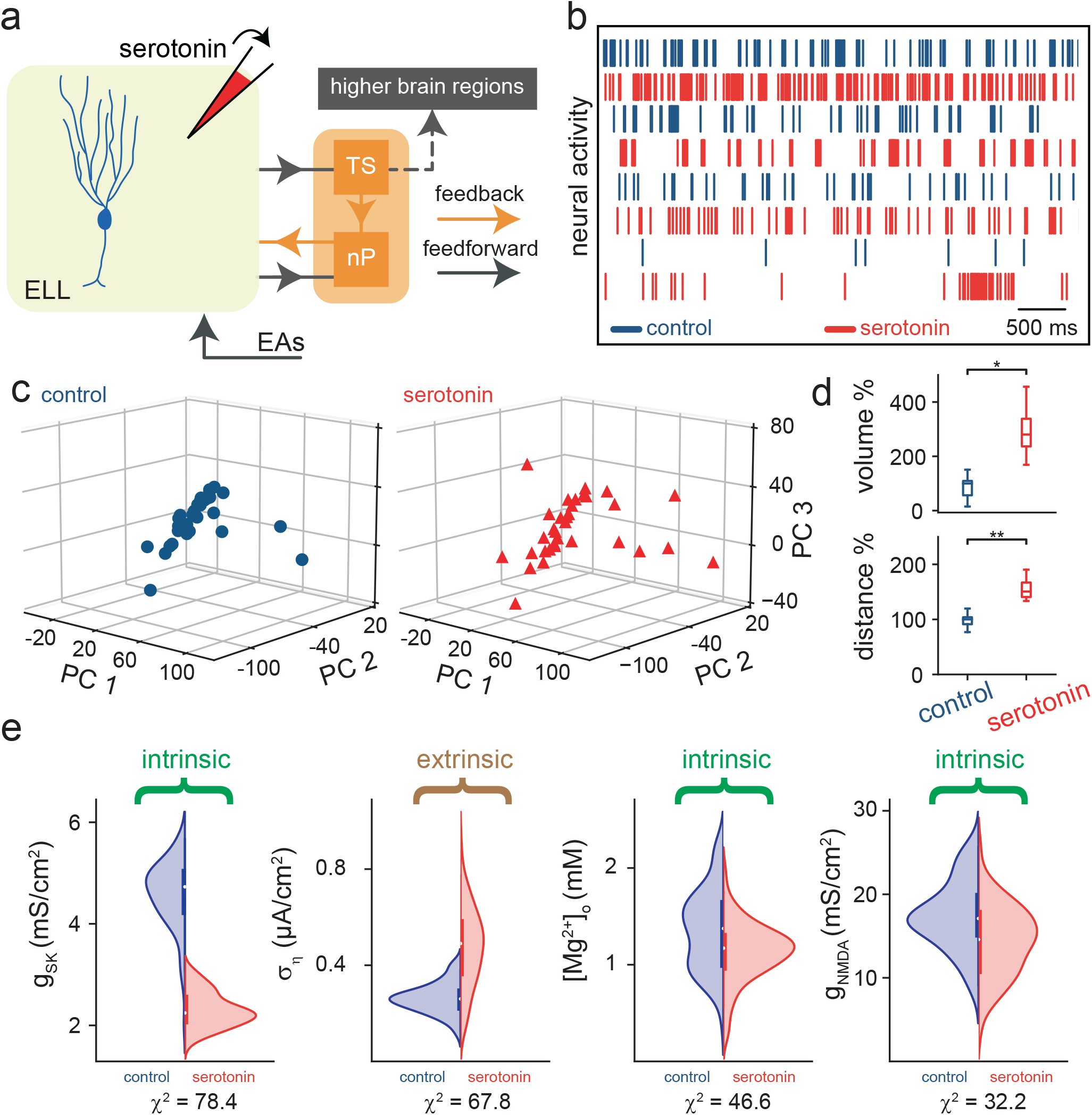
Exogenous serotonin application increases spiking heterogeneities within the ELL pyramidal cell population. **A)** Simplified diagram of feedforward (black) and feedback (orange) electrosensory pathways. Serotonin was applied exogenously in the vicinity of the ELL pyramidal cell that was recorded from using a pipette (see Methods). **B)** Spiking activities of four example ELL pyramidal cells before (dark blue) and after (red) serotonin application. **C)** Plots of our dataset using the first three principal components (i.e., PC 1, PC 2, and PC 3) that together account for 82%of the variance before (left) and after (right) serotonin application, where each datapoint represents a neuron. Notice how the datapoints occupy a significantly larger volume after serotonin application, thereby implying higher pairwise distances on average. **D)** Box plots showing that the serotonin-to-control volume (top) and pairwise distance (bottom) ratios were significantly increased (volume: *p = 0.00002, two-sample t-test; pairwise distance: **p = 0.0002, two-sample t-test). **E)** Violin plots showing distributions of pairwise distances before (blue, left) and after (orange, right) serotonin application for the four model parameters that needed to be varied significantly in order to account for changes in spiking activity seen experimentally. Consistent with previous experimental data, the value of g_SK_ (the SK maximum channel conductance) needed to be reduced the most (left). However, the value of *σ_η_* (the standard deviation of extrinsic synaptic input) needed to be also increased, explaining why the spiking activities of ELL pyramidal cells are more heterogeneous after serotonin application. The values of g_NMDA_(the NMDA receptor maximum conductance) and [Mg^2+^]_o_ (the extracellular magnesium concentration) needed to be reduced to a lesser extent.

## Discussion

### Summary of results

In order to gain insight as to how extrinsic vs. intrinsic mechanisms affect spiking heterogeneities in vivo, we used a combination of computational modeling and biophysical approaches in which we recorded from a sensory neural population under different conditions. Overall, our computational model successfully reproduced the different spiking activities in terms of mean firing rate and burst firing seen across our experimental data as quantified by differences in the ISI distribution. Further analysis revealed that parameters associated with both the intensity and mean level of extrinsic synaptic input needed to be varied the most in order to account for spiking heterogeneities seen experimentally. We then performed manipulations in order to test this prediction. First, we pharmacologically inactivated feedback pathways, which was predicted to reduce spiking heterogeneities. Analysis of experimental data obtained after inactivation revealed lower spiking heterogeneities. Further analysis of our model revealed that, in order to reproduce changes in spiking activity due to pharmacological inactivation, the values of parameters associated with both the intensity and mean level of extrinsic synaptic input needed to be reduced overall. Second, we applied the neuromodulator serotonin. Based on previous experimental results showing that such application inhibits SK currents, we initially expected a decrease in spiking heterogeneities following serotonin application. However, experimental data instead showed an increase in spiking heterogeneities. Analysis of model parameters whose values needed to be changed significantly to reproduce serotonin-dependent changes in spiking activity revealed that, while the SK conductance decreased with serotonin application, the intensity of extrinsic synaptic input also increased, thereby leading to an increase in spiking heterogeneity.

### Role of extrinsic vs. intrinsic mechanisms towards determining spiking properties

Our model makes several important predictions as to the nature of the mechanisms that mediate heterogeneous firing activities of ELL pyramidal cells in vivo and how these are affected by manipulations such as pharmacological inactivation of feedback pathways as well as serotonin application. Under control conditions, our model predicted that spiking heterogeneities were most influenced by extrinsic synaptic input, but also by differences in several intrinsic properties such as membrane maximum conductances (SK and NMDA), the concentration of IP3, and cellular morphology as quantified by the somatic-to-dendritic area ratio. Experimental data has shown wide variation in the expression of NMDA receptors (40), SK channels (39), and IP3Rs (61) across the ELL pyramidal cell population, with superficial pyramidal cells displaying the highest expression and deep pyramidal cells the lowest (see (30) for review). It is also known that the somatic-to-dendritic area ratio is highest for superficial pyramidal cells, with the largest apical dendritic trees, and lowest for deep pyramidal cells, with the smallest apical dendritic trees (27). As such, the fact that these parameters needed to be varied in order for our model to reproduce spiking heterogeneities seen in vivo suggests that graded expression of the corresponding mechanisms across the ELL pyramidal cell population contributes to spiking heterogeneities.

In order to reproduce the effects of pharmacological inactivation of feedback pathways, we needed to reduce both the intensity and the mean level of extrinsic synaptic input in the model. Although expected, this result is nevertheless important because it shows that the model is able to correctly reproduce changes in spiking activity due to pharmacological inactivation by reducing extrinsic synaptic input without any a priori information as to what the effects of the manipulation were. This is because the maximum likelihood algorithm was only presented with experimental data obtained under a given condition at any given time. This suggests that our model can correctly reproduce the effects of various factors, such as membrane conductances, calcium mobilization, and synaptic input, on spiking activity in ELL pyramidal cells. Importantly, other parameters involved in calcium mobilization (the IP3R and SRECA maximum flux rates), as well as the SK channel maximum conductance needed to be reduced. Pharmacological inactivation of feedback pathways should strongly reduce presynaptic input, and therefore lower calcium entry via NMDA receptors, an effect strongly dependent on pre-synaptic activity. The reduction in both the IP3R and SERCA maximum flux rates thus reflect changes in calcium mobilization that occur as a result of reduced calcium entry. In other words, the reduced SERCA maximum flux rate acts in part to compensate for the expected resulting decrease in internal calcium concentration. Further studies are needed to understand how pharmacological inactivation affects calcium mobilization in ELL pyramidal cells.

In order to reproduce the effects of serotonin application on spiking activity, we needed to primarily reduce the value of the SK channel maximum conductance in our model. This agrees with previous experimental data showing that serotonin application inhibits SK currents in the electrosensory system, which reduces the spike afterhyperpolarization and thereby promotes burst firing (50, 51). However, we also needed to increase the intensity of extrinsic synaptic input, which was not predictable from previous results. How does serotonin application lead to more intense synaptic input? One possibility is that activation of serotonergic pathways not only affects ELL pyramidal cells but also interneurons within the ELL (41), whose increased firing activity would be reflected in the increased noise intensity incorporated into the model. Alternatively, the reduced SK current after serotonin application can lead to increased membrane resistance, thereby increasing the voltage fluctuations due to synaptic input, which would also be reflected by an increase in synaptic input intensity. Further studies are needed to test these two hypotheses. Additionally, our model suggests that changes in firing activity due to serotonin are, to a lesser extent, due to changes in both the NMDA receptor maximum conductance (g_NMDA_) as well as the extracellular magnesium concentration [Mg^2+^]_o_. We note that previous studies in other systems have shown that serotonin downregulates NMDA-mediated currents in the prefrontal cortex (62, 63), and that magnesium interacts with the serotonergic system (64–66). As such, our modeling prediction are physiologically plausible and provide new avenues of research.

Finally, it should be noted that the experimental data considered in this study came from the lateral segment of the ELL. Previous studies have shown that the ELL consists of three parallel maps of the body surface (i.e., centro-medial, centrolateral, and lateral) with each afferent trifurcating to contact ELL pyramidal cells within each map (67, 68). Expression patterns of SK channels and NMDA receptors (39, 40), as well as serotonergic innervation (50) have also been shown to vary across segments. Therefore, future studies should test whether the model presented in this study can successfully predict the nature of the mechanisms mediating heterogeneities across ELL segments.

### Feedback pathways promote spiking heterogeneities, implications for electrosensory processing

Our results showed that spiking heterogeneities across the ELL pyramidal cell populations are primarily due to feedback input. While we only considered spiking in the absence of stimulation in the current study, it is important to realize that the trial-to-trial variability of the response to repeated stimulus presentations is determined by such activity (69, 70). As such, our analysis predicted that feedback will strongly alter the trial-to-trial variabilities of ELL pyramidal cell responses to repeated stimulus presentation. This has important implications for studying population coding (i.e., how information about sensory input is represented by neural populations in order to give rise to perception and behavior), as previous studies have shown that the so-called “noise correlations” (i.e., correlations between the trial-to-trial variabilities of neural responses) can strongly influence information transmission (see (9, 71–73) for review). In fact, previous studies have demonstrated that noise correlations in the ELL are context-dependent, which is in part due to feedback input (74, 75). Further studies are needed in order to understand how heterogeneous spiking activities across the ELL pyramidal cell contribute to population coding.

### Viability of using computational modeling in order to understand the relative contributions of extrinsic vs. intrinsic mechanisms towards spiking heterogeneity

Our study highlighted for the first time that it is feasible to use a computational model in order to reproduce the spiking activities observed across a given sensory neural population in vivo. As mentioned above, previous studies have used similar approaches, but instead focused on experimental data obtained in vitro (24, 25). Specifically, a recent modeling effort has shown that computational modeling, parameterized by in vitro data, can allow for the detection of causal relationships between transcriptomics, morphology, intrinsic membrane conductances, and electrophysiology in the mouse visual cortex (24). Another recent modeling effort has successfully shown that calcium-activated potassium channels play a critical role towards mediating burst firing in CA3 pyramidal cells (25). In the case of the electrosensory system, previous mathematical models were shown to correctly reproduce the spiking dynamics seen in vitro but not in vivo (58). This is because these did not include calcium mobilization, in particular calcium entry via NMDA receptors, which activates SK channels (56, 59). Our results showed that including calcium mobilization, as well as realistic synaptic input, was sufficient to correctly reproduce the different spiking patterns seen in vivo across ELL pyramidal cells. It is thus likely that the models used in other systems and brain areas that correctly reproduce spiking activity in vitro will need to be extended in order to reproduce spiking activity obtained under in vivo conditions.

Finally, our results demonstrating that spiking heterogeneities are primarily due to synaptic input from feedback pathways are likely to be applicable in other systems. This is because there exist many similarities between the electrosensory and mammalian systems. Specifically, ELL pyramidal cell receptive field structures and response properties are similar to those of neurons within the mammalian visual system (see (76) for review). There are also important similarities between the electrosensory and mammalian vestibular systems in terms of optimized coding of natural stimuli (77–81). This is expected given that both the electrosensory and vestibular systems have evolved from the lateral line (82).

## Conclusion

In conclusion, we have shown that it is feasible to use mathematical modeling to investigate the relative contributions of different mechanisms towards determining spiking heterogeneities in vivo. Indeed, our model made predictions that were validated experimentally. This approach is generic and thus can be generalized to other neuronal systems. Moreover, because of the important similarities between the electrosensory and other systems, it is likely that our results highlighting the role of extrinsic synaptic input in promoting spiking heterogeneities due to feedback are generally applicable.

## Materials and Methods

A detailed description of the methods is provided in SI Materials and Methods. All procedures were approved by the McGill University’s Animal Care Committee (#5285) and were in compliance with the guidelines of the Canadian Council on Animal Care. In brief, fish were purchased from suppliers and were housed according to published guidelines (83). Recordings from ELL pyramidal cells were obtained using standard techniques (84). Our dataset consisted of a total of n=243 neurons under control conditions. Of these, n=52 neurons were recorded before and after pharmacological inactivation of feedback pathways, while another n=33 neurons were recorded before and after serotonin application. Pharmacological inactivation of feedback pathways and serotonin application were also implemented using standard techniques (52, 53). Baseline activity was quantified using the interspike interval (ISI) distribution. Our model consisted of three components: (I) A two-compartmental Hodgkin-Huxley type model that describes the membrane potentials in both the soma and the dendrite. (II) A flux-balance model that describes calcium mobilization in both the cytosol and the endoplasmic reticulum (ER). (III) Realistic synaptic bombardment that mimic in vivo conditions obtained by integrating all the excitatory and inhibitory pre-synaptic inputs on the dendritic compartment (85). The model was fitted to experimental data using maximum likelihood. Model parameters that needed to be varied the most were identified by looking at distributions of pairwise distances. In the case of pharmacological inactivation of feedback pathways and serotonin application, model parameters were identified by comparing distributions obtained before and after each manipulation.

## Acknowledgments

This research was funded by the Fonds de Recherche du Québec – Nature et Technologies grant to M.J.C. and A.K; the Canadian Institutes of Health Research grant to M.J.C; the Natural Sciences and Engineering Research Council of Canada (NSERC) grant to M.J.C. and A.K.; the NSERC-CREATE Complex Dynamics award to A.A.; the Healthy Brains, Healthy Lives and the Canada First Research Excellence Fund support to A.A.

## Author Contributions

A.A., M.J.C., and A.K. designed the research. A.A., M.H., M.M.M., and S.F. performed the research. A.A. developed the computational model. A.A., M.H., and M.M.M. analyzed the data. A.A. M.J.C., and A.K. wrote the manuscript. M.J.C. and A.K. supervised and acquired the financial support for this project.

