## Supporting Information for "Decoding the relative contributions of extrinsic and intrinsic mechanisms in mediating heterogeneous spiking activities of sensory neurons in vivo using computational modeling"

**This file includes:**

1. Supporting Methods
2. Supporting Figures: S1 to S3
3. Supporting Table: S1 to S3
4. Supporting References

### Ethics statement

All animal procedures were approved by the McGill University animal care committee (# 5285) and were performed in accordance with the guidelines of the Canadian Council on Animal Care.

### Animals

Wave-type weakly electric fishes Apteronotus leptorhynchus (N=19) and Apteronotus albifrons (N=11) of either sex were used in this study. Animals were purchased from tropical fish suppliers and were housed in groups (2–10) at controlled water temperatures (26-29 °C) and conductivities (300-800 mS/cm) according to published guidelines (1).

### Surgery

Surgical procedures have been described in detail previously (2–6). Briefly, 0.1-0.5 mg of tubocurarine was injected intramuscularly to immobilize the animals for electrophysiology experiments. The animals were then transferred to an experimental tank ($30 \times 30 \times 10$ cm) that contained water from the animal's home tank and respired by a constant flow of oxygenated water through their mouth at a flow rate of approximately 10 ml/min. The head of the animal was locally anesthetized with lidocaine ointment (5%; AstraZeneca), then the skull was partly exposed, and a window of approximately (5 $mm^2$) was opened over the hindbrain.

### Recordings, pharmacological inactivation of feedback pathways, and serotonin application

Extracellular recordings were made from ELL pyramidal cells within the lateral segment in the absence of stimulation (i.e., in the presence of the animal's unmodulated EOD) with Neuropixels probes (Imec, Leuven, Belgium) as done previously (7, 8) using spikeGLX (Janelia Research Campus, Howard Hughes Medical Institute). The recordings were digitized at 30 kHz and stored on a hard drive for offline analysis. We used an automatic spike sorting algorithm followed by manual curation to sort spikes and identify single units. Specifically, we used Kilosort2 (`https://github.com/MouseLand/Kilosort2`) followed by manual curation using Phy2 (`https://github.com/cortex-lab/phy`). Well-isolated single ELL pyramidal cells were identified using various measures such as the firing rate, inter-spike-interval, spike waveform, and autocorrelograms. The sorted neural response activity was then imported into MATLAB (MathWorks Inc., Natick, MA USA) where custom code was used to analyse the data as described below. A total of n=158 neurons were recorded across N=5 Apteronotus leptorhynchus. The mean firing rate in the absence of stimulation was 16±9 Hz, which is consistent with that seen in previous studies (9).

For some recording sessions, ELL pyramidal cell activity was recorded as described above before and after pharmacological inactivation of feedback pathways. Pharmacological inactivation was achieved by injecting the sodium channel antagonist lidocaine into the nP bilaterally as done previously (10, 11). Briefly, drug application electrodes were made using single-barrel borosilicate capillary glass micropipettes (OD 1.5 mm, ID 0.86 mm, A-M Systems) and pulled by a vertical micropipette puller (Stoelting) and broken to a fine tip that was subsequently broken to attain a tip diameter of approximately 5 $\mu$m. All pharmacological injections were performed approximately 1250 – 1750 $\mu$m below the surface where the nP is located using a duration of 130 ms at ∼20 psi using a picospritzer (General Valve) as done previously (10, 11). As such, recordings from the same group of neurons could be compared before and after pharmacological inactivation of feedback pathways. Overall, a total of n=52 neurons were recorded across N=3 Apteronotus leptorhynchus. Before inactivation of feedback pathways, the mean firing rate in the absence of stimulation was 18±10 Hz, which is consistent with that seen in previous studies (9).

For other recording sessions, single unit recordings from n=17 ELL pyramidal cells in Apteronotus leptorhynchus (N=11) and n=16 ELL pyramidal cells in Apteronotus albifrons (N=11) in the absence of stimulation were made using electrodes filled with Woods Metal whose tip was plated with both gold and platinum (12). All recordings were amplified (AM systems 1700) and digitized at a 10 kHz sampling rate (CED 1401; Spike 2 Version 8.1 software; Cambridge Electronic Design). A threshold was used to identify action potential times. Previous studies have shown that electrophysiological properties of ELL pyramidal cells in both species are indistinguishable from one another (5, 6, 9, 13, 14). As such, data from both species were pooled. Each neuron was recorded from in the absence of stimulation before application of serotonin, which was applied exogenously using methodology previously described in (5, 6, 13, 15). Briefly, a double-barrel pipette with one barrel containing glutamate (1 mM) and the other serotonin (1 mM) was used. We relied on short latency excitatory responses to glutamate ejection in order to verify that the pipette was correctly within the vicinity of the neuron being recorded from. Glutamate and serotonin were both delivered using a picospritzer at 15 to 25 psi during 100 ms. As such, recordings from the same group of neurons could be compared before and after serotonin application. Overall, before application of serotonin, the mean firing rate in the absence of stimulation was 13±6 Hz, which is consistent with that seen in previous studies (9).

As there was no significant difference between the baseline activities recorded under control conditions in all three groups (p=0.01, one-way ANOVA), data was pooled for further analysis. Interspike intervals (ISIs) were obtained as the times between consecutive action potentials. As such, our dataset consisted of a total of n=243 neurons under control conditions. Of these, n=52 neurons were recorded before and after pharmacological inactivation of feedback pathways, while another n=33 neurons were recorded before and after serotonin application.

### Software

All model simulations were performed in python using Euler-Maruyama method (16) as the integration solver of the full stochastic model. The excitatory and inhibitory pre-synaptic inputs were generated using the algorithm described previously (17). Initially, we ran all simulations for a period of 120 s with an integration timestep of 0.005 ms (with a 200 kHz sampling rate). The membrane voltage trace was then downsampled to 20 kHz. For parameter optimization and fitting the model with extracellular data, we used Lmfit library in python (18). The code for generating the result and figures is available at: https://github.com/aminakhshi/spc_model.git.

### Model

To better understand how different mechanisms such as synaptic input, ion channel expression, and morphology contribute to observed heterogeneities in the spiking activities of ELL pyramidal cells in vivo, we developed a conductance-based mathematical model that consists of three components: (I) A two-compartmental Hodgkin-Huxley type model that describes the membrane potentials in both the soma and the dendrite. This component is in part inspired from previous work (19), but was extended to include key membrane conductances that regulate the spiking activity in vivo such as SK channels and NMDA receptors. (II) A flux-balance model that describes calcium mobilization in both the cytosol and the endoplasmic reticulum (ER). This component is expressed in terms of calcium fluxes that integrates firing activity of the membrane potentials with $[Ca^{2+}]_i$ transients (20). (III) Realistic synaptic bombardment that mimic in vivo conditions obtained by integrating all the excitatory and inhibitory pre-synaptic inputs on the dendritic compartment (21). The details of each component are provided below.
The two-compartmental component is based on the Hodgkin-Huxley formalism and captures the interactions between the soma and dendritic tree (3, 19). The equations describing the dynamics of membrane voltage in the soma ($V_S$) and dendritic compartments ($V_D$) are given by:

$$C_m \frac{dV_S}{dt} = I_{app} - I_{Na,S} - I_{K,S} - \frac{g_c}{\kappa}(V_S - V_D) - I_{leak} \quad (1)$$

$$C_m \frac{dV_D}{dt} = I_{syn} - I_{Na,D} - I_{K,D} - I_{SK} - I_{NMDA} - \frac{g_c}{1-\kappa}(V_D - V_S) - I_{leak}, \quad (2)$$

where $C_m$ is the membrane capacitance, $I_{Na,i}$ and $I_{K,i}$ are the fast inward sodium and the slow outward delayed rectifier potassium currents in the soma ($i = S$) and the dendrite ($i = D$), respectively, $I_{app}$ and $I_{syn}$ are the applied depolarization and synaptic currents to the somatic and dendritic compartments, respectively, $I_{SK}$ and $I_{NMDA}$ are the outward calcium-dependent small-conductance potassium and the inward NMDA calcium currents acting on the dendrite, and $I_{leak}$ is the passive leak current present in both compartments. The two compartments are linked together through a resistor, where $g_c$ is the maximum conductance and $\kappa$ is the somatic-to-dendritic area ratio. The currents $I_{Na,i}$ and $I_{K,i}$, ($i = S, D$), are necessary to generate somatic action potentials seen in vitro and the proper spike backpropagation that yields somatic depolarizing afterpotentials (DAPs) (19). The two currents $I_{NMDA}$ and $I_{SK}$ were added to the dendritic compartment because of the important role they play in regulating spiking activities of ELL pyramidal cells both in vitro (22) and in vivo (3, 23, 24). The kinetics of the ionic currents included in each compartment ($i = S, D$) are given by:

$$I_{Na,S} = g_{Na,S} m^2_{\infty,S}(1 - n_S)(V_S - V_{Na}) \quad (3)$$

$$I_{K,S} = g_{K,S} n^2_S (V_S - V_K) \quad (4)$$

$$I_{leak,S} = g_{leak}(V_S - V_{leak}) \quad (5)$$

$$I_{Na,D} = g_{Na,D} m^2_{\infty,D} h_D (V_D - V_{Na}) \quad (6)$$

$$I_{K,D} = g_{K,D} n^2_D p_D (V_D - V_K) \quad (7)$$

$$I_{SK} = g_{SK} \frac{[Ca^{2+}]^2_{i,D}}{[Ca^{2+}]^2_{i,D} + k^2_{Ca}}(V_D - V_K) \quad (8)$$

$$I_{NMDA} = g_{NMDA} B(V_D)[O](V_D - V_{Ca}) \quad (9)$$

$$I_{leak,D} = g_{leak}(V_D - V_{leak}), \quad (10)$$

where $g_{j,i}$ (j = Na,K,leak,SK,NMDA and i = S, D) are the maximum conductances, $V_j$ (j = Na,K,Ca,leak) are the Nernst potentials, $[Ca^{2+}]_i$ is the Ca concentration in the dendritic compartment ($\mu$M), $k_{Ca}$ is the half-maximum activation of the SK channel ($\mu$M), $B(V_D)$ is the magnesium block given by (25):

$$B(V_D) = \frac{1}{1 + \exp(-0.062 V_D)[Mg^{2+}]_o/3.57} \tag{11}$$

$[Mg^{2+}]_o$ is the extracellular magnesium concentration (mM), [O] is the open probability of the NMDA receptors defined by a Markov model comprised of three closed, one open and one desensitized states with transition rates $R_i$ (i = b,u,c,o,d,r) between them, $m_{\infty,i}$ (i = S, D) is the steady state activation of $I_{Na,i}$, and x ($x = n_i, h_D, p_D$; i = S, D) are the activation/inactivation gating variables whose steady states and time constants are denoted by $x_\infty$ and $\tau_x$, respectively, and whose dynamics are governed by the equation:

$$\frac{dx}{dt} = \frac{x_\infty - x}{\tau_x}. \tag{12}$$

In this model formalism, it was assumed that $I_{SK}$ and $I_{NMDA}$ are co-localized within spines. We therefore did not consider calcium diffusion within/between spines.
The NMDA receptor activation depends on the concentration of glutamate following principles of ligand-gated channels described previously (25), where the transition rates between unbound and bound states is based on the concentration of ligand. In our model, the timing of the release of glutamate follows a Poisson process formalism with firing rate $\lambda_{glu}$. The glutamate concentration following each release event was described by an alpha function (25):

$$glu = \frac{t - t_0}{\tau_1} e^{\frac{t-t_0}{\tau_1}} \tag{13}$$

where $t_0$ is the timing of glutamate release, determined by presynaptic spike times, and $\tau_1$ is the time constant of the alpha function. The parameter $\tau_1$ was chosen to be fast enough to prevent glutamate release events from overlapping.
The calcium mobilization model followed the flux-balance formalism; it describes fluxes across the cell and ER membranes in the dendritic compartment. Calcium mobilization across the cell membrane includes three fluxes through the NMDA receptors: $J_{NMDA} = \alpha I_{NMDA}$, where $\alpha = 1/(2F\bar{V}_D)$ (F is Faraday's constant and $\bar{V}_D$ is the volume of the dendritic compartment), plasma membrane calcium ATPases (PMCA) pumps: $J_{PMCA}$, and leak: $J_{INLeak}$. Calcium mobilization across the ER membrane, on the other hand, includes three fluxes through the IP3Rs: $J_{IP3R}$, SERCA pumps: $J_{SERCA}$, and leak: $J_{ERLeak}$. The Li-Rinzel model was adopted to describe IP3R kinetics (20, 26, 27). It is given by:

$$\frac{d[Ca^{2+}]_i}{dt} = f_c(J_{NMDA} - J_{PMCA} + J_{IP3R} + J_{ERleak} - J_{SERCA} - J_{INleak}) \tag{14}$$

$$\frac{d[Ca^{2+}]_{ER}}{dt} = f_{ER}\gamma(J_{SERCA} - J_{IP3R} - J_{ERleak}), \tag{15}$$

where $f_c$ ($f_{ER}$) is the fraction of free calcium concentration in the cytosolic (ER) component of the dendrite, and $\gamma$ is the volume ratio of cytosol to ER in the dendrite.
The fluxes through the PMCA and SERCA pumps are given by:

$$J_\eta = \nu_\eta \frac{[Ca^{2+}]_i^\iota}{[Ca^{2+}]_i^\iota + K_\eta^\iota}, \qquad \eta = \text{PMCA}, \text{SERCA}, \tag{16}$$

where $\iota$=2 is the Hill coefficient, $\nu_\eta$ are the maximum flux rates ($\mu$M/s), and $K_\eta$ are the half-maximum activations for calcium flux ($\mu$M). Fluxes due to leak across cell and ER membranes are given by:

$$J_{INleak} = \nu_{INleak}\left([Ca^{2+}]_o - [Ca^{2+}]_i\right) \tag{17}$$

$$J_{ERleak} = \nu_{ERleak}\left([Ca^{2+}]_{ER} - [Ca^{2+}]_i\right), \tag{18}$$

where $\nu_\xi$ ($\xi$ = INleak, ERleak) are the maximum flux rates ($\mu$M/s). Finally, flux through IP3Rs is adopted from the Li-Rinzel model (20) and is given by

$$J_{IP3R} = \nu_{IP3R} m^3_{\infty,IP3R} n^3_{\infty,IP3R} h^3_{IP3R}\left([Ca^{2+}]_{ER} - [Ca^{2+}]_i\right), \tag{19}$$

where $\nu_{IP3R}$ is the maximum flux rate ($\mu$M/s),

$$m_{\infty,IP3R} = \frac{[IP3]}{[IP3]+d_1} \quad (20)$$

$$n_{\infty,IP3R} = \frac{[Ca^{2+}]_i}{[Ca^{2+}]_i+d_5} \quad (21)$$

$$\frac{dh_{IP3R}}{dt} = \frac{h_{\infty,IP3R}-h_{IP3R}}{\tau_{h_{IP3R}}} \quad (22)$$

$$h_{\infty,IP3R} = \frac{Q_2}{Q_2+[Ca^{2+}]_i} \quad (23)$$

$$\tau_{h_{IP3R}} = \frac{1}{a(Q_2+[Ca^{2+}]_i)} \quad (24)$$

$$Q_2 = d_2\frac{[IP3]+d_1}{[IP3]+d_3}. \quad (25)$$

Note that [IP3] is the cytosolic concentration of IP3 in the dendrite ($\mu$M).

The synaptic input current, $I_{syn}$, was used to represent all the stochastic background excitatory and inhibitory pre-synaptic activity, defined by $W_x(t)$ ($x = E, I$), applied to the dendritic compartment (21, 28). It has been previously suggested that the power spectra of excitatory and inhibitory background activity in synaptic inputs exhibit a power law spectrum, $S_x(f) \sim 1/f^{\beta}$ , where $\beta$ determines the exponent of steepness of the slope in the power spectrum (21). Based on this, the total synaptic input incorporated into the model can be described by:

$$I_{syn}(t) = \sum_{x\in(E,I)} (\sigma_x \times W_x)(t) = \sigma_\eta \times W_\eta(t), \quad (26)$$

where $\sigma_\eta$ is the total noise intensity obtained by integrating the two components, $W_\eta(t)$ is the sum of both excitatory and inhibitory pre-synaptic inputs reconstructed using the following expression:

$$W_\eta(t) = \sum_{\omega} \sqrt{S_x(\omega)}\cos(\omega-\phi(\omega)), \quad (27)$$

with $\omega = 2\pi f$ and $\phi(\omega) \in [0, 2\pi]$ is a random phase multiplied by the spectrum in the frequency domain. Once the spectrum is reconstructed, the synaptic time series can be obtained using inverse Fourier transform of the spectrum as described previously (17).

### Fitting the model to experimental data

In order to reproduce the firing activities of ELL pyramidal cells obtained in vivo, we used maximum likelihood optimization to fit the interspike interval distributions generated by the computational model to those obtained experimentally. Specifically, we followed the following four steps when estimating model parameters of each cell: (I) We first chose random parameter values within the physiological range and simulated the model for a duration of 120 s. (II) We then removed the first 20 s of the simulation to exclude transient spikes, allowing us to obtain spike times by detecting the peaks of somatic membrane potentials using a threshold of 20 mV. (III) After peak-detection, we computed the ISIs between consecutive spike times from recorded data and model simulations and generated ISI distributions with 2 ms bin width. (IV) Finally, we estimated the root mean squared error (RMSE) values between ISI distributions of the model and the data and used maximum likelihood optimization (29) to update parameters in order to minimize the error with respect RMSE. RMSE was estimated within each bin of the ISI distribution according to the equation:

$$RMSE = \sqrt{\frac{1}{\Pi}\sum_{i=1}^{\Pi}(ISI_i^{model}-ISI_i^{data})^2}, \quad (28)$$

where i denotes the bin's number and $\Pi$ the total number of bins in the histogram. Tolerance was chosen in such a way that the absolute value of the difference between model outcomes and experimental data was less than 10% for each bin of the ISI distribution.

### Pairwise distance estimation

Each fitted parameter set describes the firing activity of one ELL pyramidal cell. To identify those parameters involved in generating heterogeneity between the firing properties of these cells, we first took all optimized parameter values obtained from fitting the ISI distributions and normalized them using the z-score. To remove outliers that could be caused by over fitting, we only kept parameter values within the range of $\pm 2$ from the mean after performing the z-score. Then we estimated the averaged sum of Euclidean distances of parameter values between pair of cells $D_{p_k^i}$, given by

$$D_{p_k^i} = \sum_{j \neq i}^{n} \sqrt{(p_k^i - p_k^j)^2}, \qquad i \in \{1,..,n\}, \tag{29}$$

where $p_k$ denotes each parameter in the model, whereas i and j indicate the cell number. Finally, we normalized these distances across all parameters to be between 0 and 1.

### Volume estimation

We estimated the volume of points in our principal component space by first finding the data points with minimum and maximum coordinates and divided the data points into two subsets. Then, we determined the point with maximum distance from the line that connects the first two data points to form a triangle. We repeated these steps on the two lines formed by the triangle until the convex hull contained all points in the dataset. We used convex hull algorithm from Scipy library for the volume estimation (30, 31).

### Identifying model parameters most influenced by pharmacological inactivation of feedback pathways and serotonin application

In order to identify model parameters responsible for reproducing the effects of either pharmacological feedback inactivation or serotonin application on spiking activity, we concatenated values of each parameter in the model to obtain parameter distributions before and after each manipulation. As mentioned above, the same set of neurons were recorded before and after each manipulation. We then used $\chi^2$-test to quantify the variation:

$$\chi^2 = \sum_{i} \frac{(O_i - E_i)^2}{E_i}, \tag{30}$$

where $E_i$ ($O_i$) denotes the parameter values before (after) manipulation and $i-1$ shows the degree of freedom in the $\chi^2$ estimation ($i = 10$). We counted the frequency of parameter values within each $i^{th}$ bin of the histogram for the estimation. To ensure the convergence of the estimator, bins with empty values for the control condition were ignored in the estimation.

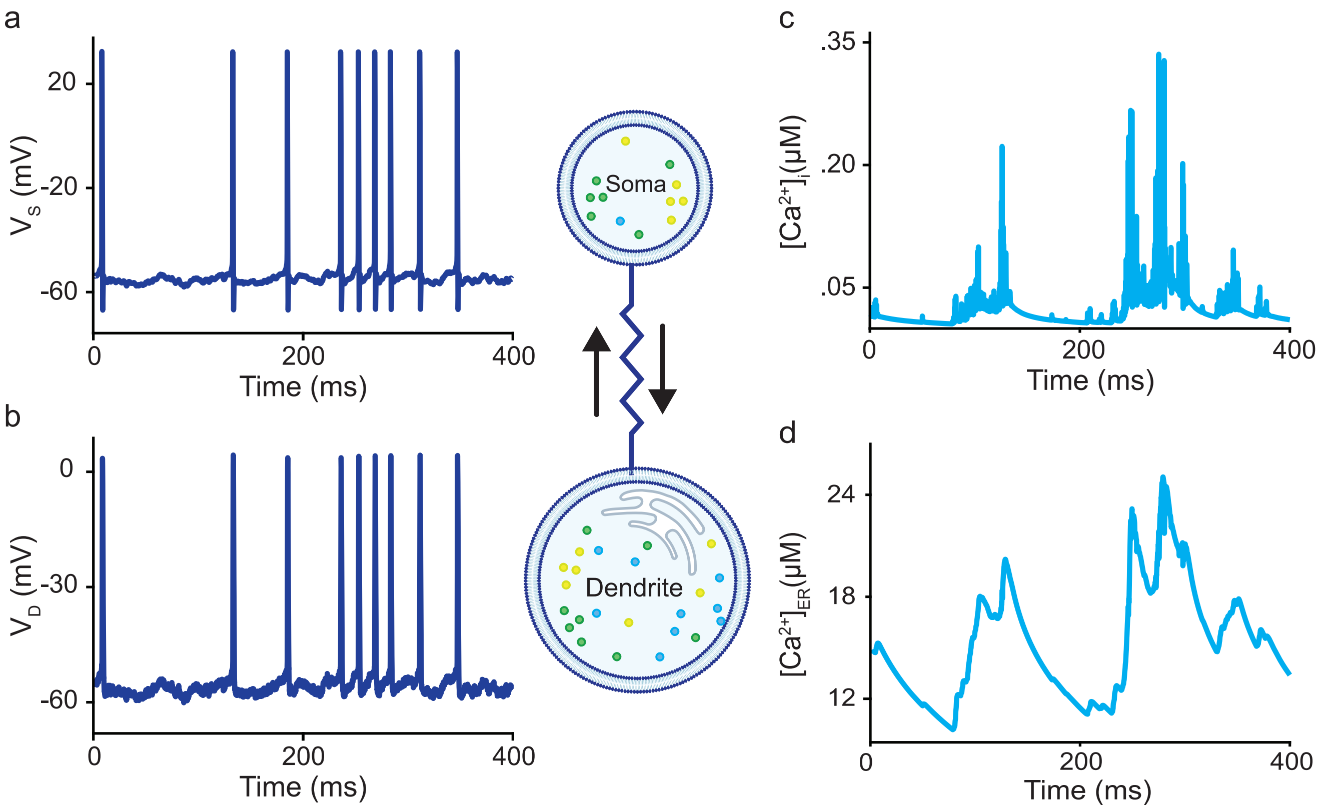


**Fig. 1. A,B)** Time series of the somatic (A) and dendritic (B) membrane voltages from the model. **C,D)** Times series of the calcium concentration within the cytosol (top) and the ER (bottom). The middle shows a simplified schematic of the model.

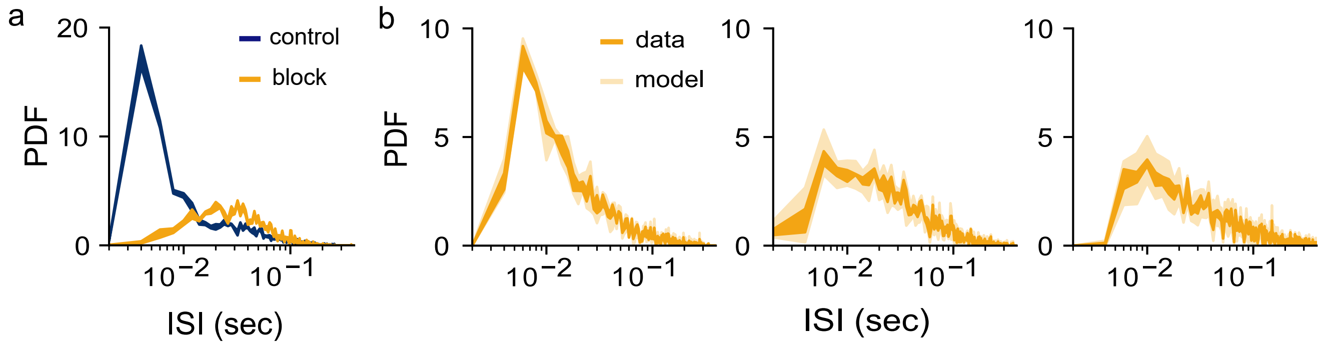


**Fig. 2. A)** ISI distributions from an example ELL pyramidal cell before (dark blue) and after (orange) pharmacological inactivation of feedback pathways. **B)** ISI distributions obtained experimentally after pharmacological inactivation (orange) and from the model (light orange) for three example ELL pyramidal cells.

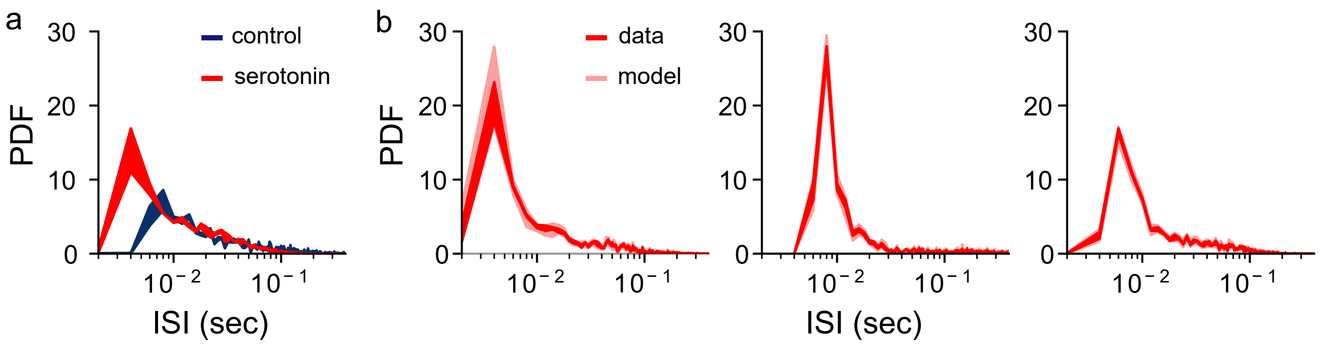


**Fig. 3. A)** ISI distributions from an example ELL pyramidal cell before (dark blue) and after (red) serotonin application. **B)** ISI distributions obtained experimentally after serotonin application (red) and from the model (light red) for three example ELL pyramidal cells.

**Table 1.** Parameter values used in the two-compartmental Hodgkin-Huxley model. Values obtained by fitting the model simulations to the data are obtained for each cell.

| Parameter | Description | Value | unit |
|---|---|---|---|
| $C_m$ | membrane capacitance | 1 | $\mu$F/cm$^2$ |
| $V_{Na}$ | $Na^+$ channel Nernst potential | 40 | mV |
| $V_K$ | $K^+$ channel Nernst potential | -88.5 | mV |
| $V_{Ca}$ | $Ca^{2+}$ channel Nernst potential | 100 | mV |
| $V_{leak}$ | leak Nernst potential | -70 | mV |
| $g_{Na}$ | $Na^+$ channel maximum conductances in the Soma/Dendrite | 55/5 | mS/cm$^2$ |
| $g_{Dr}$ | $K^+$ channel maximum conductances in the Soma/Dendrite | 20/15 | mS/cm$^2$ |
| $g_{leak}$ | Soma $\leftrightarrow$ Dendrite leak maximum conductance | 2 | mS/cm$^2$ |
| $g_{SK}$ | SK channel maximum conductance | 2 | mS/cm$^2$ |
| $g_{NMDA}$ | NMDA receptor maximum conductance | 20 | mS/cm$^2$ |
| $k_{Ca}$ | half-maximum activation of the SK channel | 0.4 | $\mu$M |
| $\tau_n$ | activation time constants of $K^+$ channel in the Soma/Dendrite | 0.39/0.9 | ms |
| $\tau_{h,p}$ | inactivation time constants of $Na^+$/$K^+$ channels in the Dendrite | 1.0/5.0 | ms |
| $V_{m,n}$ | steady-state conductance curve $V_{1/2}$ of $Na^+$/$K^+$ channel activation in the Soma | -40/-40 | mV |
| $V_{m,h}$ | steady-state conductance curve $V_{1/2}$ of $Na^+$ channel activation/inactivation in the Dendrite | -40/-52 | mV |
| $V_{n,p}$ | steady-state conductance curve $V_{1/2}$ of $K^+$ channel activation/inactivation in the Dendrite | -40/-65 | mV |
| $s_m$ | steady-state conductance curve constant of $Na^+$ channel activations in the Soma/Dendrite | 3/5 | |
| $s_n$ | steady-state conductance curve constant of $K^+$ channel activations in the Soma/Dendrite | 3/5 | |
| $s_{h,p}$ | steady-state conductance curve constant of $Na^+$/$K^+$ channel inactivations in the Dendrite | -5/-6 | |

**Table 2.** Parameter values used in the NMDA receptors Markov model. Values obtained by fitting the model simulations to the data are obtained for each cell.

| Parameter | Description | Value | unit |
|---|---|---|---|
| $[Mg^{2+}]_o$ | extracellular $Mg^{2+}$ concentration | 1.0 | mM |
| $R_{b,u}$ | binding/unbinding rates between the closed ($C_0 \leftrightarrow C_1 \leftrightarrow C_2$) states | 40/12.9 | 1/s |
| $R_{o,c}$ | binding/unbinding rates between the open $\leftrightarrow$ closed ($C_2 \leftrightarrow C_o$) states | 46.5/73.8 | 1/s |
| $R_{r,d}$ | binding/unbinding rates between the desensitized $\leftrightarrow$ closed ($C_2 \leftrightarrow C_d$) states | 6.8/8.4 | 1/s |

**Table 3.** Parameter values used in the flux-balance Calcium model. Values obtained by fitting the model simulations to the data are obtained for each cell

| Parameter | Description | Value | unit |
|---|---|---|---|
| $\nu_{PMCA,SERCA}$ | maximum flux rates through PMCA/SERCA pumps | 30/22.5 | $\mu$M/s |
| $K_{PMCA,SERCA}$ | half-maximum activation for calcium fluxes through PMCA/SERCA | 0.45/0.105 | $\mu$M |
| $\nu_{INleak,ERleak}$ | maximum flux rates through cell and ER membranes | 0.05/0.03 | $\mu$M/s |
| $\nu_{IP3R}$ | maximum flux rate of IP3R | 30 | $\mu$M/s |
| $d_{1,3}$ | dissociation constant of IP3 | 0.13/0.9434 | $\mu$M |
| $d_{2,5}$ | dissociation constant of $Ca^{2+}$ inhibition/activation | 1.049/0.08234 | $\mu$M |
| a | binding constant of $Ca^{2+}$ inhibition | 9 | 1/s |
| [IP3] | cytosolic concentration of IP3 in the dendrite | 0.3 | $\mu$M |
| $f_{c,ER}$ | fraction of free calcium concentration in the cytosolic/ER components | 0.05/0.0025 | |
| $\gamma$ | volume ratio of cytosol to ER in the dendrite | 9 | |
